# IL-15 re-programming compensates for NK cell mitochondrial dysfunction in HIV-1 infection

**DOI:** 10.1101/811117

**Authors:** Elia Moreno Cubero, Stefan Balint, Aljawharah Alrubayyi, Ane Ogbe, Rebecca Matthews, Fiona Burns, Sarah Rowland-Jones, Persephone Borrow, Anna Schurich, Michael Dustin, Dimitra Peppa

## Abstract

Dynamic regulation of cellular metabolism is important for maintaining homeostasis and can directly influence immune cell function and differentiation including Natural Killer (NK) cell responses. Persistent HIV-1 infection leads to a state of chronic activation, subset redistribution and progressive NK cell dysregulation. In this study we examined the metabolic processes that characterise NK cell subsets in HIV-1 infection, including adaptive NK cell subpopulations expressing the activating receptor NKG2C, which expand during chronic infection. These adaptive NK cells exhibit an enhanced metabolic profile in human cytomegalovirus (HCMV) infected HIV-1 seronegative individuals. However, the bioenergetic advantage of adaptive CD57+NKG2C+ NK cells is diminished during chronic HIV-1 infection, where NK cells uniformly display reduced oxidative phosphorylation (OXPHOS) and limited fuel flexibility upon CD16 stimulation. Defective OXPHOS was accompanied by increased mitochondrial depolarisation and structural alterations indicative of mitochondrial dysfunction, suggesting that mitochondrial defects are restricting the metabolic plasticity of NK cell subsets in HIV-1 infection. The metabolic requirement for receptor stimulation was alleviated upon IL-15 pre-treatment enhancing mammalian target of rapamycin complex1 (mTORC1) activity and NK cell functionality in HIV-1 infection, representing an effective strategy for pharmacologically boosting NK cell responses.

## Introduction

Natural Killer (NK) cells play a key role in antiviral immunity and a number of studies implicate NK cells as critical contributors to immune control of HIV-1 (Fauci et al., 2005) (Martin et al., 2002) (Martin et al., 2007). It is well documented that persistent HIV-1 infection alters the subset distribution and functional capacity of NK cells (Peppa et al., 2018) (Bradley et al., 2018) (Altfeld et al., 2011). In particular HCMV co-infection, a hallmark of HIV-1 infected cohorts, triggers a dramatic expansion of differentiated NK cells expressing the activating receptor NKG2C, which prototypically characterises adaptive NK cells (Guma et al., 2006) (Mela and Goodier, 2007) (Brunetta et al., 2010). These expanded NKG2C+ NK cells co-express CD57, a marker of maturation, and are influenced by HLA-E presented peptides (Hammer et al., 2018) (Rolle et al., 2018). Notably they resemble memory CD8 T cells and are imbued with enhanced capacity for antibody dependent cellular cytotoxicity (ADCC), in particular increased IFN-γ production attributed to epigenetic remodelling of the IFNG locus (Schlums et al., 2015) (Lee et al., 2015). The adaptive reconfiguration of NK cells during HIV-1 infection is further delineated by loss of the transcription factor promyelocytic leukaemia zinc finger protein (PLZF), and downstream signalling molecules such as FcεRI-γ (Peppa et al., 2018), which partly overlap with NKG2C expression. While these attributes are considered important in the functional specialisation of these NK cells, the development of these features under conditions of continuous stimulation/persistent inflammation during HIV-1 infection could lead to the establishment of functionally and metabolically exhausted NK cells akin to exhausted CD8 T cells. In keeping with this notion, chronic stimulation of adaptive NK cells through NKG2C ligation was recently found to lead to a molecular programme of exhaustion that is shared between NK cells and CD8 T cells (Merino et al., 2019). Exhausted CD8 T cells are characterised by a number of metabolic defects, however, to date it remains unexplored how persistent HIV-1 infection contributes to the metabolic remodelling of NK cell subsets. NK cell exhaustion could influence responses to CD16 engagement linked to vaccine-induced protective immunity against HIV-1 infection and phenotypes of viral control (Scully and Alter, 2016). Such knowledge represents a critical first step in our understanding of the metabolic machinery of NK cell subsets with distinct immunologic features that can be potentially targeted for developing new or complementary antiviral approaches aimed at enhancing ADCC responses.

Whereas metabolic regulation of T cell function is well characterised, evidence of the importance of immunometabolism in facilitating robust NK cell functions is gradually emerging. Notably, impaired NK cell cellular metabolism has been implicated in obesity and cancer, providing a new framework for understanding NK cell functionality (Michelet et al., 2018) (Cong et al., 2018). Studies from both human and murine models have demonstrated that NK cells activated through cytokine stimulation exhibit substantial increases in the rates of both glycolysis and oxidative phosphorylation (OXPHOS) pathways (Marcais et al., 2014) (Donnelly et al., 2014) (Keating et al., 2016; O’Brien and Finlay, 2019). However, the metabolic requirement of NK cells for optimal effector function, in terms of IFN-γ production, depends on the specific activation stimuli. In particular, receptor mediated activation through anti-NK1.1 and anti-Ly49D in mice appears to require OXPHOS as an essential second signal for IFN-γ production and appears more susceptible to metabolic inhibition compared to cytokine stimulation (Keppel et al., 2015).

Different metabolic programmes are adopted by different NK cell subsets and recent evidence suggests that distinct metabolic fingerprints underline the fate of NK cells (Pfeifer et al., 2018; Schafer et al., 2019). Along these lines adaptive NK cells emerging in response to HCMV infection in HIV-1 seronegative adults have been reported to engage different metabolic pathways exhibiting enhanced mitochondrial fitness relative to their canonical counterparts (Cichocki et al., 2018). These adaptive CD57+NKG2C+ NK cell subpopulations display superior respiratory capacity supported by increased mitochondrial mass, mitochondrial membrane potential and upregulated genes involved in the electron transport chain (ETC). The enhanced mitochondrial metabolism in adaptive NK cell subsets was found to be regulated by the chromatin-modifying transcriptional regulator AT-rich interactive domain-containing protein 5B (ARID5B) (Cichocki et al., 2018). These features may explain the increased functionality of adaptive NK cell subsets in HCMV seropositive HIV-1 seronegative donors, especially augmented IFN-γ production through CD16 dependent recognition. The latter has been linked to increased activity of the mammalian target of rapamycin complex1 (mTORC1) pathway, a well-established regulator of multiple metabolic processes (Schlums et al., 2015). However, knowledge of NK cell metabolism in other chronic viral infections such as HIV-1, where HCMV coinfection is almost universal and plays a defining role in shaping the NK cell pool, remains in its infancy.

In this study we sought to address the metabolic features of NK cell subsets during chronic HIV-1/HCMV co-infection. Our findings show that inefficient OXPHOS and loss of mitochondrial integrity in NK cells during HIV-1 infection influences responses to activating CD16 receptor stimulation. Such defects could be overcome *in vitro* by IL-15 metabolic reprogramming reducing the dependence for receptor-mediated metabolic remodelling.

## Results

### NK cells from treatment naïve viraemic HIV-1 infected patients exhibit reduced oxidative metabolism

To study the metabolic profile of NK cells in HIV-1 infection, accounting for the influence of HCMV co-infection, we isolated NK cells from HCMV seropositive treatment-naïve HIV-1 positive donors and HCMV seropositive HIV-1 negative donors (referred to as controls (CTR)). All donors in this study were HCMV+ and screened for the presence of adaptive CD57+NKG2C+ NK cells (**Fig. S1**) as previously described (Hammer and Romagnani, 2017). An extracellular flux analyser was utilised to analyse their metabolic requirements. The glycolytic rate assay was used to estimate the extracellular acidification rate (ECAR), a measure of lactate production and anaerobic glycolysis at the basal state and after the addition of rotenone and antimycin A, which interfere with complex I and complex III of the electron transport chain (ETC), respectively and 2-deoxy-glucose (2-DG) a competitive inhibitor of glycolysis. Consistent with previous reports baseline activity of NK cells was low in both groups and a non-significant trend towards lower basal ECAR was noted in the HIV infected group (**Fig. 1A, B**). No significant differences in the rates of NK cell glycolysis or glycolytic reserve were detected between the two groups, indicating that there were no major modulations of glycolytic pathways at baseline (**Fig. 1C, D, E**). Next, we assessed mitochondrial respiration by measuring oxygen consumption rates (OCR), a measure of OXPHOS, at basal levels and following the addition of a stressor mix including oligomycin (an ATP synthesis inhibitor), carbonyl cyanide-4 (trifluoromethoxy) phenylhydrazone (FCCP; uncoupling synthesis of ATP from the ETC), and rotenone and antimycin A at the indicated time points (**Fig. 1F**). At rest NK cells utilise preferentially OXPHOS and although no difference in the basal OCR levels was observed in the two groups (**Fig. 1G**) both maximal respiration rate and spare respiratory capacity (SRC) were significantly reduced in NK cells from HIV-1 infected donors (**Fig. 1H, I**) suggestive of altered oxidative metabolism.

**Fig. 1.**
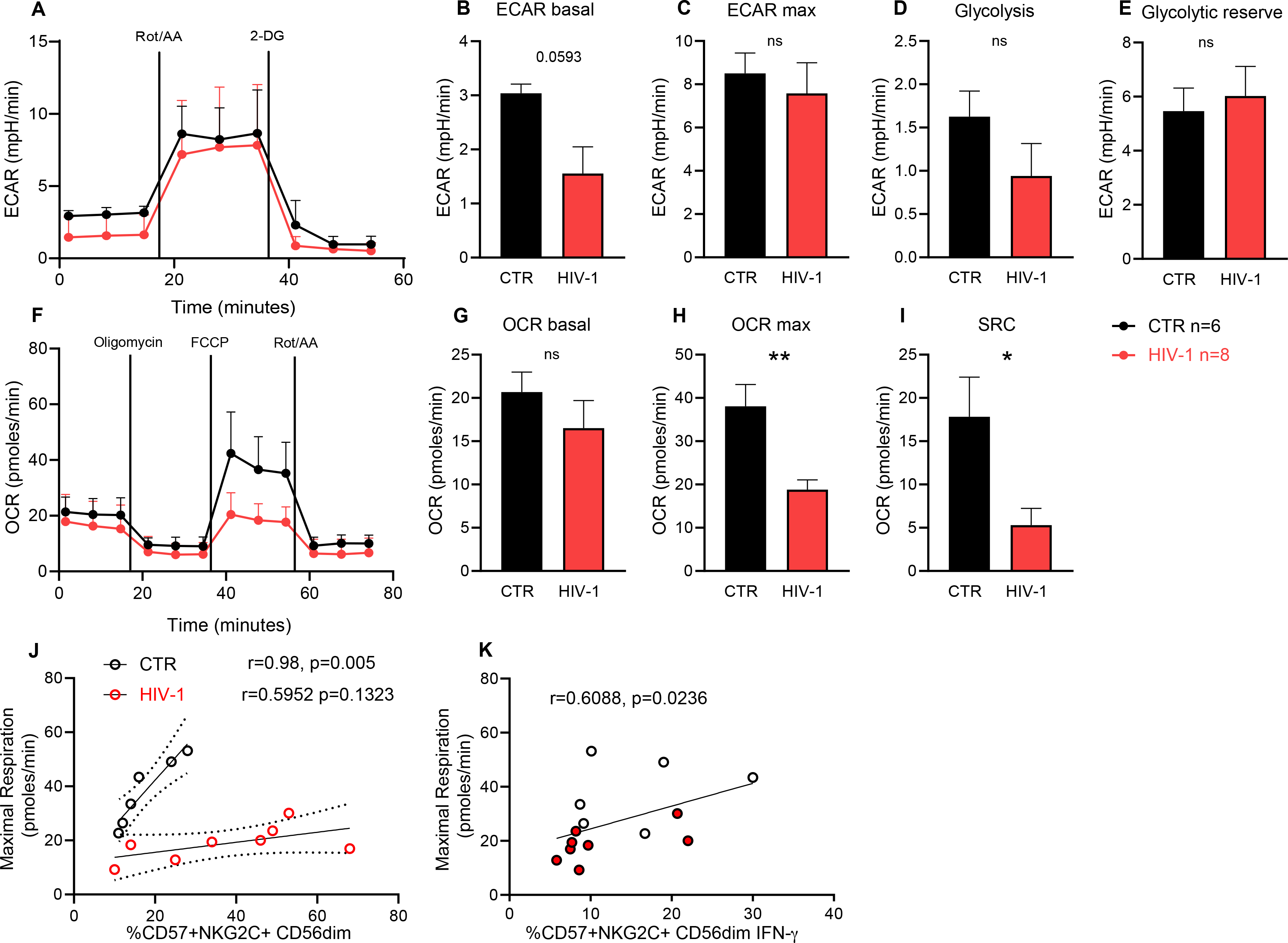
Analysis of the basal metabolic profile of NK cell in control (CTR) and HIV-1 infected donors. **(A)** Real-time analysis of aerobic glycolysis (ECAR) in purified NK cells **(B)** Basal ECAR, **(C)** maximal ECAR, **(D)** glycolysis and (**E)** glycolytic reserve in isolated NK cells from n=6 controls (CTR) and n=8 chronic HIV-1 patients (HIV-1). **(F)** Real-time analysis of oxygen consumption rate (OCR) in isolated NK cells from both groups of study. (**G)** Basal OCR, **(H)** maximal OCR (OCR max) and **(I)** Spare respiratory capacity (SRC). Bars show mean ± SEM. Correlation between maximal OCR and **(J)** *ex vivo* percentage of adaptive CD57+NKG2C+ CD56dim NK cells and **(K)** proportion of CD57+NKG2C+CD56dimIFN-γ+ after CD16 triggering. The non-parametric Spearman test was used for correlation analysis. Sample triplicates were used for Seahorse assays. Significance determined by Mann-Whitney *U* test, ns: non-significant, *p<0.05, **p<0.01

Given that adaptive CD56dimCD57+NKG2C+ NK cells are expanded in HIV-1 infection, as a consequence of more frequent HCMV reactivations and ongoing inflammation, representing a large proportion of the peripheral NK cell pool (**Fig. S1**), we sought to determine the relationship between the magnitude of these cells and respiratory capacity. Whereas a positive correlation was observed between the frequencies of adaptive NK cells and maximal respiration in HCMV+ controls (CTR) (**Fig. 1J**) as previously reported (Cichocki et al., 2018), this association was not observed in HIV-1 infected donors, suggesting that the superior metabolic profile of NKG2C+ NK cells is diminished in HIV-1 infection (**Fig. 1J**). Adaptive NK cells are less sensitive to innate cytokines but display augmented responses and in particular higher IFN-γ production through CD16 activation (Schlums et al., 2015). The relationship between the metabolic fitness and effector response in adaptive NK cells upon CD16 triggering in the cohort as a whole is reflected in the correlation between IFN-γ production and maximal respiration (**Fig. 1K**).

### OXPHOS is required for NK cell IFN-γ production following CD16 crosslinking

To investigate the requirement of OXPHOS for receptor mediated activation of NK cells we tested the ability of CD56dim NK cells to produce IFN-γ following anti-CD16 stimulation in the presence or absence of oligomycin. Addition of oligomycin significantly impaired NK cell IFN-γ production in response to CD16 triggering in the control group (**Fig. 2A, B**). However, this effect was less prominent in CD56dim and NK cell subsets in HIV-1 infected individuals in keeping with an already suppressed capacity to utilise mitochondrial OXPHOS (**Fig. 2C**, **Fig. S2A, C, D, F)**). Of note the presence of oligomycin did not affect NK cell survival at the concentration used during the 6 hour stimulation and no decrease in the levels of CD16 expression with oligomycin were observed (data not shown). Glycolytic inhibition by adding the inhibitor 2-DG in the assay abrogated IFN-γ production by NK cells in both study groups (**Fig. 2A-C & Fig. S2A-F**). Inhibition of pyruvate transport into the mitochondria with UK5099 had a more pronounced effect on IFN-γ production by CD56dim and adaptive NK cells in HIV-1 infection as compared to HCMV+ controls (**Fig. 2A, C & Fig. S2D, F**). Inhibition of glutamine, an alternative OXPHOS fuel, with BPTES had a significant effect in reducing IFN-γ from adaptive NK cells in HIV-1 infection only (**Fig. S2D,F**). Pre-treatment with etomoxir blocking fatty acid transport into the mitochondria, had no obvious effect on IFN-γ production by CD16 activated NK cells in both the control and HIV-1 infected group, suggesting that fatty acids are not utilised by NK cells under these conditions. (**Fig. 2A-C**) (**Fig. S2A-F**). Together these data suggest that glucose is the primary fuel for OXPHOS required for receptor stimulation by NK cells. However, in contrast to controls, in HIV-1 infection NK cells, and especially adaptive subpopulations, are less able to mobilise additional energy resources to power OXPHOS.

**Fig. 2.**
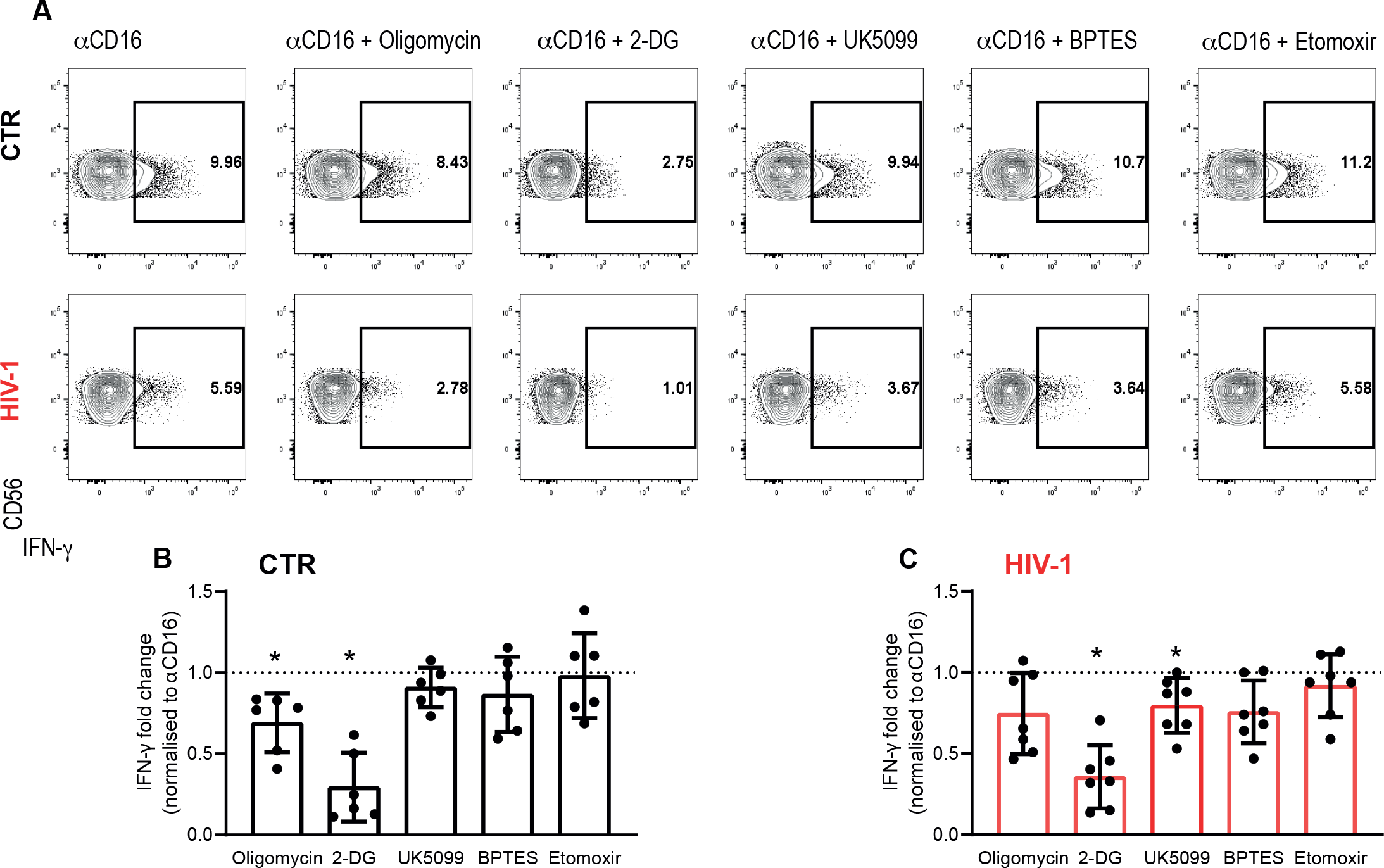
Metabolic requirements of NK cells for IFN-γ production after CD16 activation. **(A)** Representative flow plots and summary data of fold change in IFN-γ production by CD56dim NK cells following anti-CD16 plate bound stimulation from **(B)** n=6 healthy controls (CTR) and **(C)** n=7 chronic HIV-1 infected patients (HIV-1) in the presence or absence of the different metabolic inhibitors: Oligomycin (100 nM), 2-DG (50 mM), UK5099 (2 μM), BPTES (3 μM) or Etomoxir (4 μM). Fold change was normalised to anti-CD16 stimulation alone (set to 1 and indicated as a dotted line). Bars show mean ± SD and each symbol represents data from an individual donor. Wilcoxon matched-pairs signed rank test,*p<0.05.

### Adaptive NK cells in HIV-1 infection display evidence of dysfunctional mitochondria

The decrease in SRC in NK cells from HIV-1 positive donors suggested a reduced ability of these cells to produce energy in response to stress or increased work demands and prompted us to further investigate the mitochondrial health of NK cells. In particular, greater SRC is shown to correspond to enhanced mitochondrial fitness, a feature of adaptive NK cells and memory CD8 T cells (Cichocki et al., 2018) (van der Windt et al., 2012). We therefore utilised the ratiometric fluorescent dye JC-1 to assess further the mitochondrial membrane potential (ΔΨm) of NK cells, which represents a key readout of mitochondrial function. NK cells from HIV-1 infected donors displayed a lower ΔΨm especially within adaptive subsets compared to NK cells from HIV-1 seronegative donors (**Fig. 3A,B**). This is exemplified by lower red/green fluorescence intensity ratio of JC1, which does not depend on mitochondrial shape, size or density, suggesting that mitochondrial dysfunction in NK cells, especially of adaptive subpopulations which are enriched in HIV-1 positive donors, constitutes a rate limiting step in their capacity to fuel their energy demands.

**Fig. 3.**
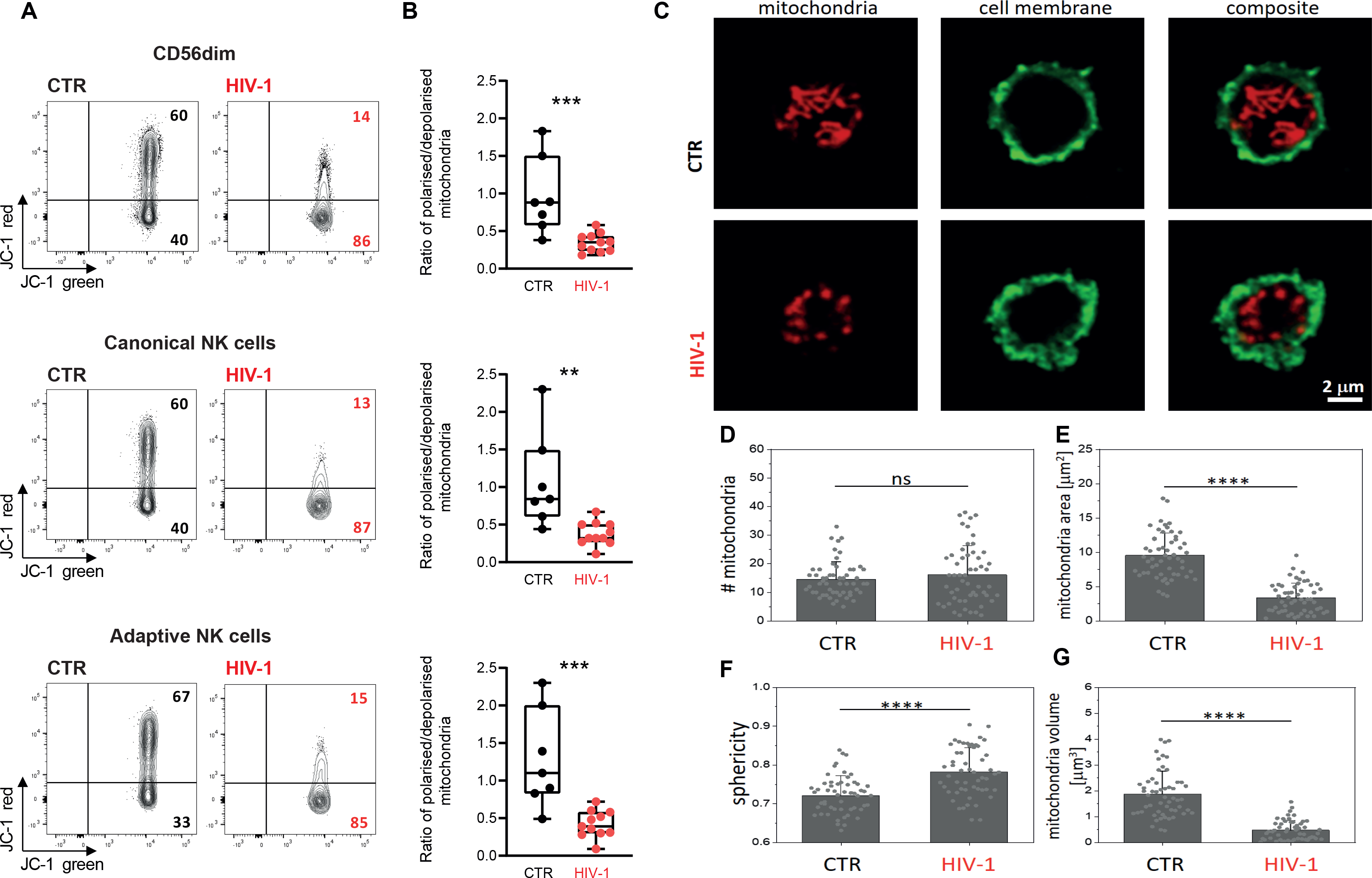
Evaluation of NK cell mitochondrial health. **(A)** Representative flow plots and (**B)** box and whisker plots depicting the ratio of polarised over depolarised mitochondrial (mitochondrial membrane potential (ΔΨm), of total CD56dim NK cells and NK cell subsets by JC-1 staining in control (CTR) and HIV-1 infected subjects (HIV-1). JC-1 red staining designates polarised mitochondria, while loss of JC-1 red shows depolarisation. The box-and-whisker plots show the median, quartiles, and range. Data from an individual subject are represented by each symbol. Significance determined by Mann-Whitney *U* test. **(C)** Representative confocal images of mitochondria (red) from purified NK cells from a control and an HIV-1 infected donor. The NK cell membrane was visualised by staining with wheat germ agglutinin (green). (**D)** Quantification of mitochondria numbers, **(E)** area, **(F)** sphericity and (**G)** volume in purified NK cells from both n=3 controls (CTR) and n=3 HIV-1 seropositive donors (HIV-1). Each symbol represents the mean value from one cell. Data are from a minimum of 50 cells from three independent donors. Bars represent mean ± SD. ns: non-significant, **p<0.01, ***p<0.001, ****p<0.0001, one-way analysis of variance (ANOVA) with Tukey’s post-hoc test.

Mitochondria are dynamic organelles that configure their morphology and number to regulate their function and distribution (Cogliati et al., 2016). To interrogate further the relationship between mitochondrial dysfunction and mitochondrial morphological changes we performed confocal imaging. Isolated NK cells from three HCMV+ HIV-1 seronegative controls and three HIV-1/HCMV seropositive donors with evidence of depolarised mitochondria were stained with MitoTracker Deep Red, that accumulates in active mitochondria and enables localisation. Our results demonstrated that despite no difference in the number of mitochondria (**Fig. 3C, D**), NK cells from HIV-1 infected individuals exhibited mitochondrial structural differences adopting a change in their configuration from a long tubular shape into shrunk small spherical shapes (**Fig. 3C, E, F, G**) and (**S3 and S4 movies**). These data suggest that loss of ΔΨm in NK cells from HIV-1 infected individuals most likely triggers a mitochondrial transformation and collapse of structural support mechanisms (Miyazono et al., 2018). Structural changes in the mitochondria have been related to bioenergetic insufficiency and increased mitochondrial sphericity in T cells in the absence of co-stimulation (Klein Geltink et al., 2017) and could also be a reflection of oxidative damage in the cell (Ahmad et al., 2013). However, analysis of reactive oxygen species (ROS) levels in NK cells from patients with chronic HIV-1 infection compared with those of NK cells from HCMV+ HIV seronegative controls did not show a significant increase in the levels of ROS in HIV-1 infection (data not shown). This could be due to a rescue mechanism, with adaptive NK cells previously described to display elevated levels of BCL-2, which may be important in limiting oxidative stress (Zhang et al., 2013) (Cichocki et al., 2018).

The enhanced oxidative metabolism observed in adaptive NK cells from HCMV+ HIV-1 seronegative donors has been reported to depend partly on increased expression of *ARID5B* and induction of genes encoding components of the ETC, including *UQCRB,* the ETC complex III gene (Cichocki et al., 2018). To determine whether there were any differences in the expression of *ARID5B* in isolated NK cells from HCMV+ controls and HIV-1 infected individuals we performed quantitative RT-PCR (qRT-PCR). We confirmed increased levels of *ARID5B* expression in NK cells from HIV-1 infected donors, in keeping with greater expansions of adaptive NK cells in HIV-1 infection, relative to controls (**Fig. S5A**). Our findings therefore suggest that mitochondrial structural disorganisation rather than decreased expression of ARID5B could be potentially affecting the performance of the ETC and metabolic adaptation of NK cells in HIV-1 infection.

### Treatment with IL-15 compensates for the metabolic defects of NK cells in HIV-1 infection

IL-15 plays a vital role in NK cell survival and differentiation and in priming NK cells for enhanced effector function *in vivo* (Cooper et al., 2002). We therefore investigated whether IL-15 priming *in vitro* could improve NK cell responses to stimulation via CD16 by circumventing the metabolic requirements for efficient receptor mediated activation in HIV-1 seropositive donors. Pre-treatment with IL-15 for 48-72 hours followed by anti-CD16 stimulation led to a striking increase in IFN-γ production by total CD56dim NK cells (**Fig. 4A**) but also canonical and adaptive subsets (**Fig. S6A-D**). Inhibition of OXPHOS with oligomycin in primed NK cells did not affect IFN-γ production in response to receptor stimulation while a small difference in the presence of 2-DG persisted (**Fig. 4A, B**).

**Fig. 4.**
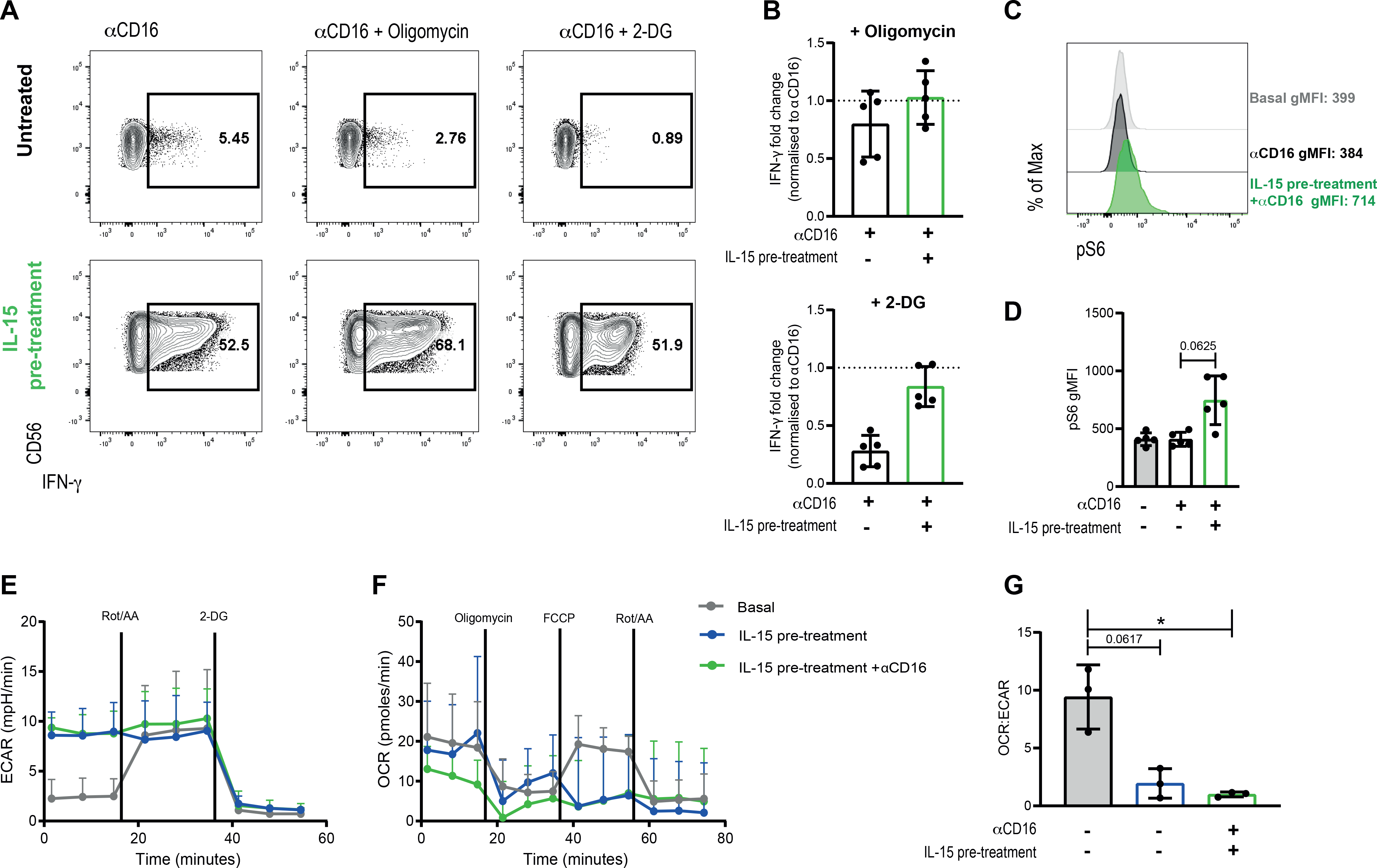
IL-15 treatment ameliorates NK cell metabolic requirements for receptor stimulation in HIV-1 infection. **(A)** Representative flow plots and (**B)** summary data of fold change in IFN-γ production by CD56dim NK cells following stimulation with anti-CD16 with or without IL-15 pre-treatment and in the presence or absence of the following inhibitors: Oligomycin (100 nM) or 2-DG (50 mM), from n=5 HIV-1 infected donors. Data normalised to anti-CD16 stimulation alone, set to 1 and indicated as a dotted line in the graph. Bars mean ± SD and each symbol indicates individual responses. **(C)** Representative histograms and **(D)** summary bar charts of pS6 expression levels to evaluate mTORC1 activity after stimulation in CD56dim NK cells from n=5 HIV-1 infected individuals. Statistics by Wilcoxon matched-pairs test. (**E)** Real-time analysis of aerobic glycolysis (ECAR), **(F)** oxygen consumption (OCR) and **(G)** basal OCR:ECAR ratio in purified NK cells from n=3 HIV-1 donors in the presence or absence of anti-CD16 and/or IL-15 pre-treatment. *p<0.05 by paired Student’s *t*-test.

IL-15 is known to stimulate multifaceted metabolic activities during maturation of NK cells via the mammalian target of rapamycin complex 1 (mTORC1), a key regulator of cellular metabolism (Marcais et al., 2014). IL-15 signalling and activation of NK cells leads to enhanced glucose uptake and enhanced functionality. Consistent with this IL-15 priming of NK cells from HIV-1 infected donors prior to anti-CD16 stimulation induced greater phosphorylation of the translational regulator S6 kinase (S6), consistent with enhanced mTORC1 signalling in CD56dim NK cells (**Fig. 4C, D**) and subsets (data not shown).

To further evaluate the metabolic effects of IL-15 pre-treatment in the presence or absence of anti-CD16 stimulation in isolated NK cells from HIV-1 infected patients we performed extracellular flux assays to directly analyse metabolic responses. NK cells treated with IL-15 displayed a higher basal ECAR especially in the presence of CD16 coated beads relative to resting NK cells (**Fig. 4E**). No significant increase in the cellular respiratory function as measured by OCR was observed either with IL-15 pre-treatment alone or with CD16 stimulation (**Fig. 4F**). A decreased dependence on OXPHOS was observed in IL-15-treated NK cells especially in the presence of CD16 triggering as evidenced by the decreased ratio of OCR:ECAR compared to basal levels (**Fig. 4G**). These findings indicate that IL-15 increased NK cell metabolism, specifically glycolysis, by-passing the “OXPHOS requirement” for receptor stimulated IFN-γ production.

## Discussion

Burgeoning evidence indicates that metabolic plasticity is important for fine tuning immune cell function. Whereas most studies have focused on T cell metabolism during human chronic viral infections, much less is known about the metabolic lifestyle of NK cells in this setting. In this study we assessed for the first time the metabolic profile of NK cell subsets and requirements for production of IFN-γ following receptor stimulation, via CD16 crosslinking, during chronic HIV-1 infection. Our results implicate dysregulated metabolism as an upstream driver of NK cell functional fate. NK cells in viraemic HIV-1 infection are characterised by impaired oxidative phosphorylation, reduced metabolic reserve and mitochondrial defects. IL-15 pre-treatment effectively eliminated the metabolic requisite for receptor stimulation boosting NK cell responses.

In general, resting NK cells are quiescent as evidenced by the low basal levels of glycolysis and OXPHOS. However, NK cells in chronic HIV-1 infection display an inability to rely on mitochondrial function under conditions of increased energy demands as evidenced by the reduction in both maximal respiration and SRC. Although we did not formally test the metabolic profile of adaptive CD57+NKG2C NK cells in this study, due to sample limitations and potential effect of sorting on the redox state of the cells (Llufrio et al., 2018), these cells are enriched and represent a sizeable proportion of the peripheral NK cell pool in HIV-1 infection. Notably, there was no correlation between CD57+NKG2C+ NK cells from HIV-1 infected donors and maximal respiration. This suggests that NK cells in HIV-1 infection do not acquire the characteristic metabolic profile defining HCMV adaptive (CD57+NKG2C+) NK cells in HIV-1 seronegative individuals with NK cell memory-like increased OXPHOS (Cichocki et al., 2018). Alternatively, adaptive subpopulations expressing NKG2C could gradually lose their bioenergetic advantage with progressive HIV-1 infection as a consequence of persistent activation and evolving dysregulation.

Upon stimulation NK cells configure their metabolic machinery to meet their energy demands and synthesize molecules required for NK cell effector function. Depending on the activating stimulus and length of stimulation NK cells can up-regulate glycolysis and OXPHOS. In this study we focused on receptor stimulation via CD16, a potent activator of NK cell function and an important means of NK cell antiviral control in HIV-1 infection. Global OXPHOS inhibition significantly reduced receptor stimulated NK cell IFN-γ production, which was further attenuated by inhibition of glycolysis in both HIV-1 infected and control individuals, in keeping with glucose being the predominant fuel driving OXPHOS. Similar effects were observed in murine models utilising NK1.1 and Ly49D activating receptor stimulation (Keppel et al., 2015), supporting the notion that OXPHOS is a metabolically required second signal for IFN-γ production mediated via ITAM coupled adapters. Inhibition of alternative fuels, such as glutamine and fatty acids, did not significantly affect IFN-γ production by NK cells in HCMV+ control donors, while an effect was observed for NK cell subsets in HIV-1 infection for glutamine and pyruvate suggesting a loss of metabolic plasticity.

Metabolic inflexibility has been recently described to underline dysfunctional HIV-specific CD8 T cells (Angin, 2019). In particular, the polyfunctionality of CD8 T cells from spontaneous HIV-1 controllers was more reliant on mitochondrial function rather than glycolysis and differences in glucose dependency described in SIV-specific CD8 T cells have been found to correlate with their capacity to supress SIV infection. We therefore reasoned that the observed inability of NK cells to utilise OXPHOS during HIV-1 infection could reflect dynamic changes in the mitochondria, which are essential hubs of metabolic activity determining the function and fate of immune cells. A global mitochondrial defect was supported by decreased mitochondrial membrane potential (Δψm) in NK cells in HIV-1 infection. Notably, isolated NK cells from HIV-1 infected patients displayed marked changes in mitochondrial architecture characterised by the presence of small spherical mitochondria. The shape of the mitochondria may directly impact their bioenergetic function, with elongated mitochondria being associated with more efficient ATP production, and mitochondrial dysfunction resulting in fragmentation described in a number of pathological conditions (Chan, 2019). Interestingly the delicate balance between mitochondrial fusion and fission controls mitochondrial shape, number and size and is important for function (respiratory capacity) and quality control of mitochondria, raising the possibility that alterations in mitochondrial dynamics in HIV-1 infection could influence NK cell metabolic reprogramming. Along these lines mitochondrial fusion regulated by optic atrophy-1 (OPA-1) was found to facilitate tight cristae organisation in memory T cells (Tm) cells, with continued entrance of pyruvate into mitochondria and efficient ETC activity (Buck et al., 2016). In contrast, effector T cells with fissed mitochondria displayed diffuse internal structures and less efficient OXPHOS, and pharmacological enforcement of mitochondrial fusion promoted oxidative capacity and superior functionality (Buck et al., 2016). It would therefore be important to dissect in future studies the intricate relationship between the modulation of mitochondrial shape and NK cell energetic state and downstream cellular processes, such as mitophagy (O’Sullivan et al., 2015), providing a framework to study these crucial biological interactions on metabolic regulation of NK cell subsets during chronic viral infections.

The observed alterations in the metabolic machinery of NK cells during chronic HIV-1 infection influencing their mitochondrial integrity, bears similarities to exhausted CD8 T cells. In particular exhausted CD8 T cells during chronic viral infections show extensive mitochondrial alterations with an impaired ability to use mitochondrial energy supply coupled to the programmed death-1 (PD-1) pathway (Schurich et al., 2016) (Fisicaro et al., 2017) (Bengsch et al., 2016). Signalling via PD-1 promoting exhaustion of CD8 T cells has been found not only to affect mitochondrial morphology but also to represses the key metabolic transcriptional regulator peroxisome proliferator-activated receptor-γ (PPAR-γ) coactivator 1a (PGC-1a), controlling energy metabolism and mitochondrial biogenesis, suppressing OXPHOS and glycolysis in virus specific T cells (Bengsch et al., 2016). Upregulation of inhibitory molecules on NK cells could also serve as a pathway supressing their metabolic fitness. Recently chronic stimulation of adaptive NKG2C+ NK cells *in vitro* led to a marked induction of checkpoint inhibitory receptors PD-1 and lymphocyte activation gene-3 (LAG-3) sharing epigenetically driven programmes with exhausted CD8 T cells (Merino et al., 2019). Moreover, a role for LAG-3 in regulating CD4 T cell metabolism and mitochondrial biogenesis has been identified (Previte et al., 2019). We have previously reported a modest increase in the levels of PD-1 expression on NK cells in HIV-1 infection (Peppa et al., 2018), however, the role of inhibitory receptors in controlling NK cell metabolism remains to be further defined.

Pre-treatment of NK cells with IL-15 alleviated the need for OXPHOS for CD16 receptor stimulated IFN-γ production extending observations from murine models (Keppel et al., 2015). Our results implicate IL-15 increased mTORC1 activity, an important regulator of NK cell metabolism, in upregulating glycolysis. The remaining dependence on glucose could reflect an overall NK cell increased dependency to glycolysis or a switch to a post-transcriptional regulatory mechanism for IFN-γ production as described for T cells (Chang et al., 2013). Notably, the effects of IL-15 priming on NK cell metabolism and functionality have important implications for clinical translation in HIV-1 infection. The beneficial effects of IL-15 immunotherapy and IL-15 superagonist ALT-803, capable of activating both NK cells and CD8 T cells, have been highlighted *in vivo* in non-human primate models (Webb et al., 2018) and such an approach is currently being tested in a phase I clinical trial to facilitate clearance of latent HIV-1 reservoirs (NCT02191098). The ability of IL-15 to enhance ADCC and augment NK cell mediated killing of HIV-infected target cells *ex vivo* (Garrido et al., 2018) may prove vital in the development of a functional cure for HIV. Our findings together with a recent report of IL-15 mediated metabolic reprogramming improving the efficacy of HIV-specific CD8 T cells from non-controllers (Angin, 2019), highlight the complementary effects of such an approach to simultaneously re-invigorate multiple arms of the immune response.

In summary, our results demonstrate that NK cells in HIV-1 infection exhibit metabolic defects affecting both canonical and adaptive subsets underlined by mitochondrial dysfunction/structural alterations and bearing similarities to exhausted CD8 T cells described in chronic viral infections. These findings identify mitochondrial centred dysfunction as a key area for future research and as a promising target for future combined reconstitution therapies. IL-15 pre-treatment bypasses the metabolic requirement for OXPHOS upon CD16 mediated activation and potentiates NK cell function, which is of considerable clinical interest in the field of HIV as a powerful multipronged therapeutic strategy. Further insights into the potential role of inhibitory receptors will provide us with a more comprehensive understanding of the integration of signals and the role of metabolism in fine tuning the range and potency of NK cell antiviral functions.

## Material and Methods

### Study Subjects

Cryopreserved peripheral blood mononuclear cells (PBMCs) from eleven HIV-1 infected HCMV seropositive treatment naïve patients were utilised (mean age= 38.49, range= 27-49; mean LogVL=4.89, range=4.14-6.33; mean CD4 count 475 cells/mL, range=61-720). Participants were recruited at the Mortimer Market Centre for Sexual Health and HIV Research and the Royal Free Hospital (London, UK) following written informed consent as part of a study approved by the local ethics board committee. Seven demographically age, sex and lifestyle matched HCMV seropositive HIV-1 seronegative donors (mean age=36.9, range=26-49) with sizeable adaptive NK cell responses (>10%) were used for comparison (**Fig. S1**), from whom blood was taken and cryopreserved for later use with written informed consent in accordance with the Declaration of Helsinki. All study participants were anti-Hepatitis C virus antibody negative and anti-HBsAg negative and HCMV seropositive. HCMV serology was determined at University College London Hospital clinical virology lab by the ARCHITECT CMV IgG assay (AU/mL) (Abbott Diagnostics, IL, USA).

### Flow cytometric analysis

The following fluorochrome-conjugated antibodies were used in this study: CD16 BB700, CD56 BV605, CD57 BV421 (BD Biosciences), CD3 BV650, CD14 BV510, CD19 BV510, CD56 PE Dazzle, CD16 PerCP (BioLegend), NKG2C PE and NKG2C AlexaFluor700 (R&D systems) for surface antigens; IFN-γ PE-CY7, (BD Pharmingen) and pS6 (S235/236) AlexaFluor647 (Cell Signaling) for intracellular antigens. Briefly, cryopreserved PBMC isolated from HIV-1-infected patients and healthy controls were washed in PBS, and surface stained at 4°C for 20 min with saturating concentrations of different combinations of antibodies in the presence of fixable live/dead stain (Invitrogen). Cells were then fixed and permeabilized for detection of intracellular antigens. Assessment of mitochondrial membrane polarisation and ROS production was performed by incubation with 2 μM JC-1 or 5 μM MitoSOX (Invitrogen) respectively according to manufacturer’s instructions. Samples were acquired on a BD Fortessa X20 using BD FACSDiva8.0 (BD Bioscience) and data analysed using FlowJo 10 (TreeStar). **Fig. S1** includes the gating strategy for the identification of NK cell subsets into adaptive and canonical on the basis of expression of CD57 and NKG2C.

### NK cell isolation

NK cells were enriched from PBMCs using a negative-selection magnetic bead kit (Miltenyi Biotec) as per the manufacturer’s instructions (>94% purity and viability).

### Extracellular flux assays

Extracellular flux assays were performed using an XFp Analyzer (Agilent technologies). Purified NK cells were seeded at 2×10^5^ cells per well in Seahorse cell plates coated with CellTak (Corning). Mitochondrial OXPHOS was evaluated using the mitostress test kit and glycolysis using the glycolytic rate assay kit (Agilent technologies). The oxygen consumption rate (OCR) and extracellular acidification rate (ECAR) were measured in XF RPMI medium supplemented with 10 mM glucose, 1 mM pyruvate and 2 mM glutamine in response to oligomycin (1 μM), FCCP (1 μM), rotenone/antimycin A (0.5 μM) and 2-DG (50 mM) (Agilent technologies). To evaluate the effect on the metabolic profile of IL-15 pre-treatment, NK cells were purified and cultured with 100 ng/mL of IL-15 for 48h, then split and further stimulated with CD16-coated beads (Milteny) or left in IL-15 supplemented media. Maximum respiration is the average OCR values post-FCCP injection. The spare respiratory capacity (SRC) was calculated as average post-FCCP injection minus basal OCR average. Glycolysis was calculated as average basal ECAR values minus post-2-DG injection values. Glycolytic reserve was calculated as average post-rotenone/antimycin A injection ECAR values minus basal ECAR values.

### Functional assays

For activation via CD16 cross-linking, 96-well flat-bottom plates (Nunc) were coated with 5 μg/mL anti-human CD16 (clone 3G8, BD Biosciences) or an isotype-matched control antibody (mIgG1κ, BD Biosciences) overnight at 4°C. Plates were washed with sterile PBS before addition of 4 × 10^5^ PBMC per well. To assess fuel flexibility cells were incubated for 6 h in the presence or absence of 100 nM Oligomycin, 50 mM 2-DG, 2 μM UK5099, 3 μM BPTES and 4 μM Etomoxir (Sigma). GolgiStop (containing Monensin, 1/1,500 concentration) (BD Biosciences) and GolgiPlug (containing brefeldin A, 1/1,000 final concentration) (BD Biosciences) were added for the last 5 h of culture. Following incubation cells were washed and stained for extracellular receptors before permeabilization and intracellular staining for IFN-γ and pS6. Where indicated cells were pre-treated with 100 ng/mL IL-15 (Miltenyi) for 48-72h.

### Confocal microscopy

Confocal imaging was performed with an Olympus FV1200 inverted microscope (Olympus) equipped with a 60× 1.4 NA oil-immersion objective at room temperature.

Prior to confocal imaging, purified NK cells were incubated with 50 nM MitoTracker Deep Red FM (ThermoFisher Scientific) for 60 min at 37°C to visualise the mitochondria. After incubation, the cells were plated onto 0.01 % Poly-L-Lysine coated 8-well IBIDI chamber for 15 min followed by fixation with 4% PFA/PBS for additional 30 min at room temperature. An additional labelling with 10 μg/mL WGA-AlexaFluor488 (ThermoFisher Scientific) for 10 min to visualize the cell membrane was performed. Images were taken with a 250 nm z-stepping. Post processing and segmentation (TrakEM2 plugin) of fluorescence images was done with ImageJ (National Institute of Health). The number of mitochondria, surface area, mitochondria volume and sphericity were obtained by 3D surface rendering of confocal images in Imaris version 9.3 (Bitplane).

### Quantification of gene expression

Total NK cells were lysed following isolation using MACS beads (Miltenyi). Total RNA was extracted using Rneasy micro kit (Qiagen) following the manufacturer’s instructions. cDNA synthesis was performed using the high capacity cDNA reverse transcriptase kit (Applied Biosystems). Real time PCR was performed using 5ng of CDNA, fast SYBR green master mix (Applied Biosystems) and the following primers: GAPDH forward 5’-CCTGCACCACCAACTGCTTA-3’; reverse 5’- GGCCATCCACAGTCTTCTGAG-3’, RPLP0 forward: 5’-GCAATGTTGCCAGTGTCTG-3’; reverse 5’-GCCTTGACCTTTTCAGCAA-3’, and ARIDB5 forward 5’-CTGAGCCTCTCCCAGCAGCA-3’; reverse 5’-CGCCTCCTCTGCCACCTTCT-3’ on a 7500 fast real-time PCR system (Applied Biosystem). GAPDH and RPLP0 were used as housekeeping genes for normalisation. Data were analysed using ΔCt method relative to housekeeping genes and for both control and HIV-1 infected subjects and expressed as log2 expression value.

### Statistics

Prism 8 (GraphPad Software) and OriginPro 9.1 (OriginLab) were used for statistical analysis as follows: the Mann–Whitney *U*-test or Student’s *t*-test were used for single comparisons of independent groups, the Wilcoxon-test or Student’s paired *t*-test were used to compare two paired groups. The non-parametric Spearman test was used for correlation analysis. To analyse the confocal imaging results, samples were tested for normality with a Kolmogorov–Smirnov test. The statistical significance for multiple comparisons was assessed with one-way analysis of variance (ANOVA) with Tukey’s post hoc test (\**p* < 0.05, \*\**p* < 0.01, \*\*\**p* < 0.001, and \*\*\*\**p* < 0.0001).

## Author Contributions

EMC performed experiments, contributed to study design, acquisition of data, analysis, and drafting of the manuscript; SB, AA, AO performed experiments and contributed to acquisition of data; RM, FB, SRJ, PB and MD contributed to study design, data interpretation and critical editing of the manuscript. DP: conception and design of study, data analysis and interpretation, critical revision of the manuscript and study supervision.

## Supplementary material

**Fig. S1. Gating strategy to distinguish canonical and adaptive NK cells. (A)** Representative example gated on live CD3-, CD14-, CD19- lymphocytes; CD56 and CD16 are used to identify natural killer (NK) cell subsets. **(B)** Representative examples from a healthy control and an HIV-1 infected patient showing frequencies of canonical NK cells defined as CD56dimCD57+NKG2C- and adaptive NK cells defined as CD56dimCD57+NKG2C+.

**Fig. S2. Metabolic requirements of different NK cell subsets for IFN-γ production after CD16 activation.** Representative flow plots of CD57+NKG2C- CD56dim (canonical) **(A)** and CD57+NKG2C+ CD56dim (adaptive) NK cells from a control (CTR) and an HIV-1 infected donor (HIV-1) **(D)** showing IFN-γ production following anti-CD16 stimulation in the absence and presence of the indicated metabolic inhibitors. Summary bar charts of fold change in IFN-γ production by canonical NK cells following anti-CD16 plate bound stimulation from **(B)** n=6 control individuals and **(C)** n=7 HIV-1 infected donors and by adaptive NK cells from **(E)** n=6 control individuals and **(F)** n=7 HIV-1 infected subjects in the presence or absence of the different metabolic inhibitors: Oligomycin (100 nM), 2-DG (50 mM), UK5099 (2 μM), BPTES (3 μM) or Etomoxir (4 μM). Fold change was normalised to anti-CD16 stimulation alone (set to 1 and indicated as a dotted line). Bars show mean ± SD and each symbol represents data from an individual donor. Wilcoxon matched-pairs signed rank test *p<0.05

**Movie S3.** Representative confocal z-stack, 3D projection and reconstruction of mitochondrial distribution in a primary human NK cell isolated from a CMV+ control. Mitochondria, red; cell membrane, green.

**Movie S4:** Representative confocal z-stack, 3D projection and reconstruction of mitochondrial distribution in a primary human NK cell isolated from an HIV-1 infected donor. Mitochondria, red; cell membrane, green.

**Fig. S5.** Quantification of gene expression by RT-qPCR for **(A)** *ARID5B* in isolated NK cells from n=4 HCMV+ control donors and n=4 HIV-1 infected subjects. All relative expression values were calculated by normalising against GAPDH as a housekeeping gene. **p<0.01

**Fig. S6. IL-15 treatment ameliorates NK cell subsets metabolic requirements for receptor stimulation in HIV-1 infection.** Representative flow plots **(A, B)** and summary bar charts of fold change in IFN-γ production by **(C)** canonical and **(D)** adaptive NK cells following stimulation with anti-CD16 with or without IL-15 pre-treatment and in the presence or absence of the following inhibitors: Oligomycin (100 nM) or 2-DG (50 mM), from n=5 HIV-1 infected donors. Fold change was normalised to anti-CD16 stimulation alone (set to 1 and indicated as a dotted line). Bars show mean ± SD and each symbol represents data from an individual donor. Statistics by Wilcoxon matched-pairs test.

