## Supplementary material for "IL-15 re-programming compensates for NK cell mitochondrial dysfunction in HIV-1 infection": Fig. S1

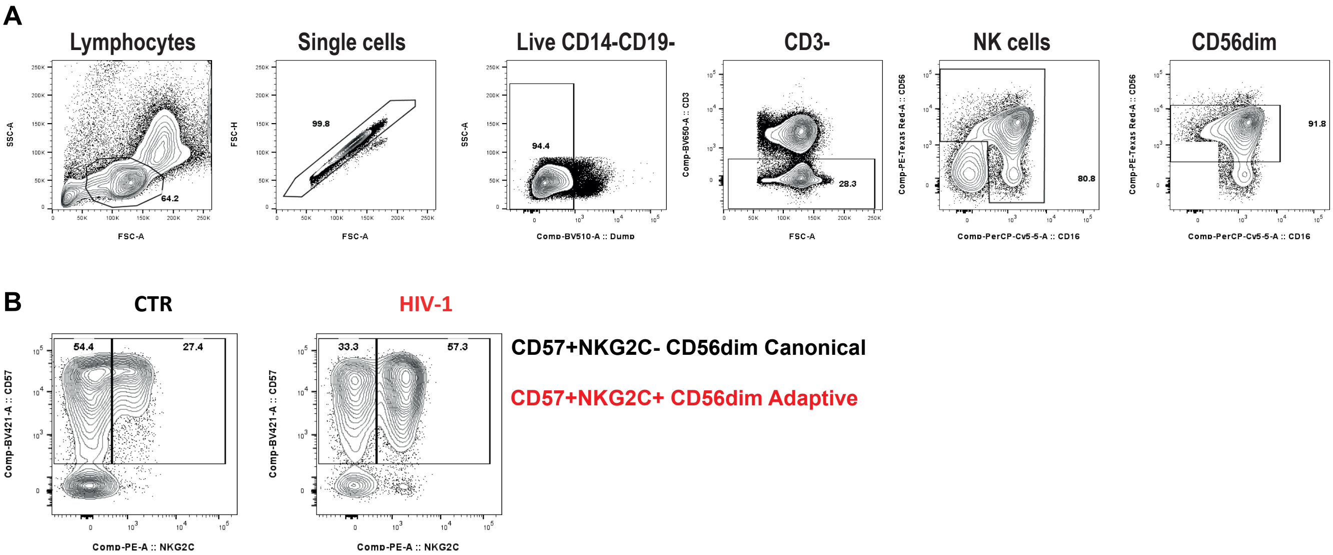


**Fig. S1. Gating strategy to distinguish canonical and adaptive NK cells. (A)** Representative example gated on live CD3-, CD14-, CD19- lymphocytes; CD56 and CD16 are used to identify Natural Killer (NK) cell subsets. **(B)** Representative examples from a healthy control and an HIV-1 infected patient showing frequencies of canonical NK cells defined as CD56dimCD57+NKG2C- and adaptive NK cells defined as CD56dimCD57+NKG2C+.
