## Supplementary material for "IL-15 re-programming compensates for NK cell mitochondrial dysfunction in HIV-1 infection": Fig. S2

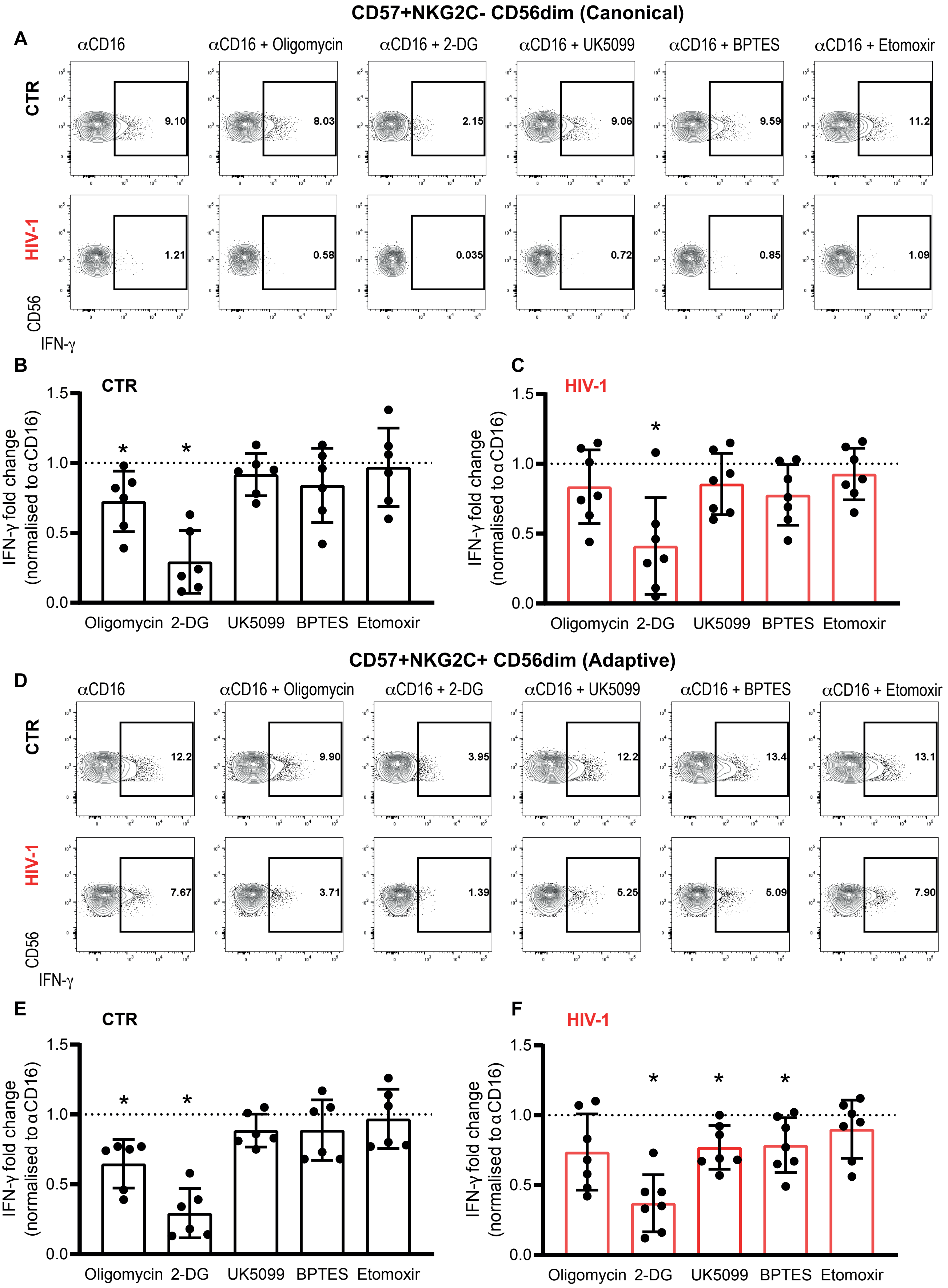


**Fig. S2. Metabolic requirements of different NK cell subsets for IFN-γ production after CD16 activation.** Representative flow plots of CD57+NKG2C- CD56dim (canonical) **(A)** and CD57+NKG2C+ CD56dim (adaptive) NK cells from a control (CTR) and an HIV-1 infected donor (HIV-1) **(D)** showing IFN-γ production following anti-CD16 stimulation in the absence and presence of the indicated metabolic inhibitors. Summary bar charts of fold change in IFN-γ production by canonical NK cells following anti-CD16 plate bound stimulation from **(B)** n=6 control individuals and **(C)** n=7 HIV-1 infected donors and by adaptive NK cells from **(E)** n=6 control individuals (CTR) and (F) n=7 HIV-1 infected subjects (HIV-1) in the presence or absence of the different metabolic inhibitors: Oligomycin (100 nM), 2-DG (50 mM), UK5099 (2 μM), BPTES (3 μM) or Etomoxir (4 μM)**.** Fold change was normalised to anti-CD16 stimulation alone (set to 1 and indicated as a dotted line). Bars show mean ± SD and each symbol represents data from an individual donor. Wilcoxon matched-pairs signed rank test *$p<0.05$.
