## Supplementary material for "IL-15 re-programming compensates for NK cell mitochondrial dysfunction in HIV-1 infection": Fig. S5

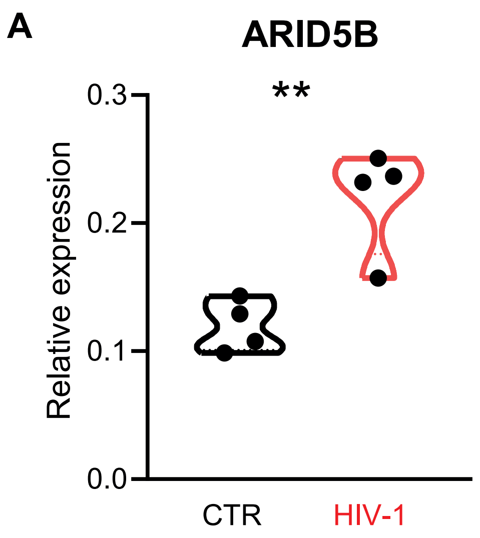


**Fig. S5.** Quantification of gene expression by RT-qPCR for **(A)** ARID5B in isolated NK cells from n=4 HCMV+ control donors (CTR) and n=4 HIV-1 infected subjects (HIV-1). All relative expression values were calculated by normalising against GAPDH as a housekeeping gene. **p<0.01
