## Supplementary material for "IL-15 re-programming compensates for NK cell mitochondrial dysfunction in HIV-1 infection": Fig. S6

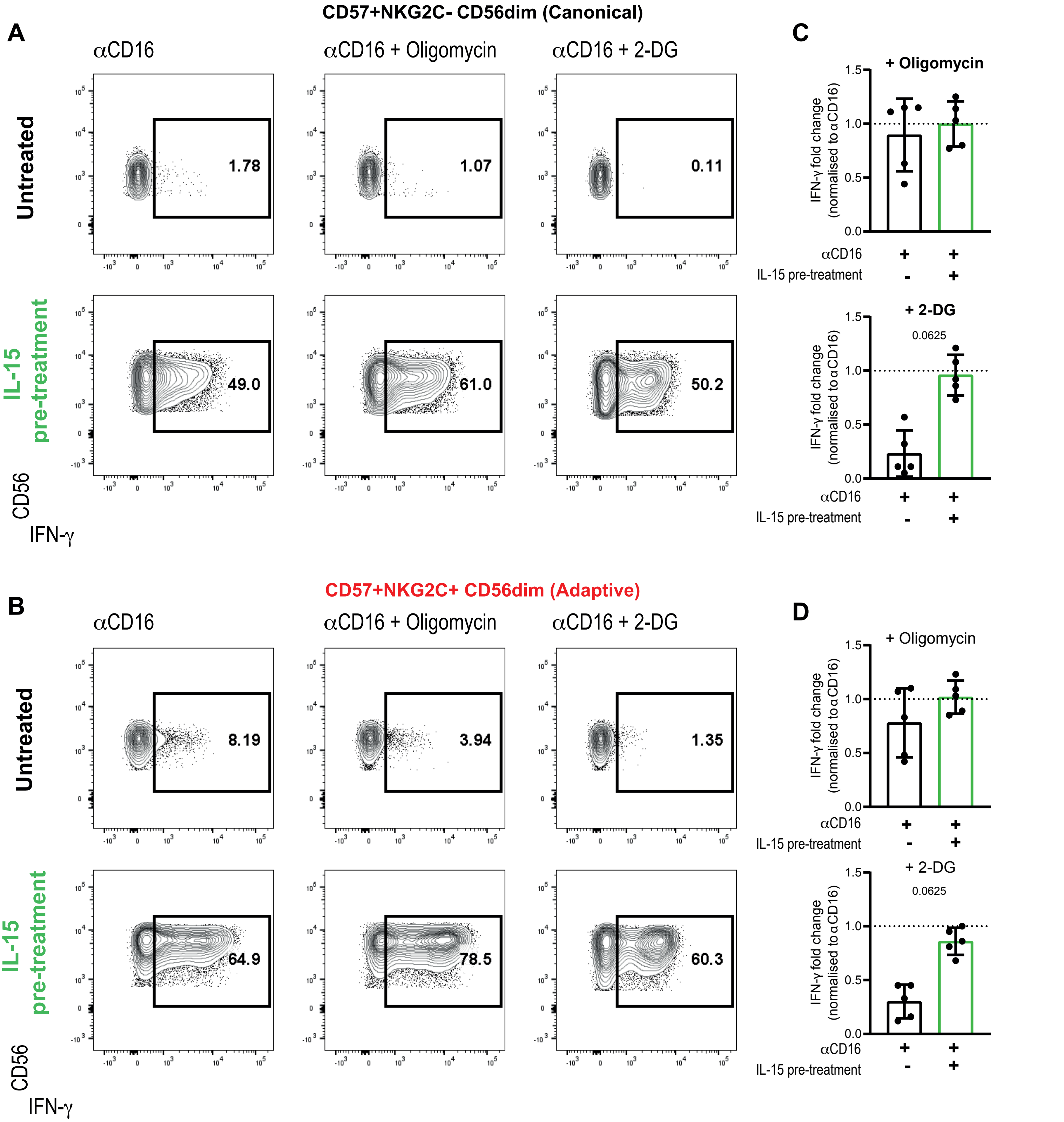


**Fig. S6. IL-15 treatment ameliorates NK cell subsets metabolic requirements for receptor stimulation in HIV-1 infection.** Representative flow plots **(A, B)** and summary bar charts of fold change in IFN-γ production by **(C)** canonical and **(D)** adaptive NK cells following stimulation with anti-CD16 with or without IL-15 pre-treatment and in the presence or absence of the following inhibitors: Oligomycin (100 nM) or 2-DG (50 mM), from n=5 HIV-1 infected donors. Fold change was normalised to anti-CD16 stimulation alone (set to 1 and indicated as a dotted line). Bars show mean ± SD and each symbol represents data from an individual donor. Statistics by Wilcoxon matched-pairs test.
